# DDIEM: Drug Database for Inborn Errors of Metabolism

**DOI:** 10.1101/2020.01.08.897223

**Authors:** Marwa Abdelhakim, Eunice McMurray, Ali Raza Syed, Senay Kafkas, Allan Anthony Kamau, Paul N Schofield, Robert Hoehndorf

## Abstract

**Background:** Inborn errors of metabolism (IEM) represent a subclass of rare inherited diseases caused by a wide range of defects in metabolic enzymes or their regulation. Of over a thousand characterized IEMs, only about half are understood at the molecular level, and overall the development of treatment and management strategies has proved challenging. An overview of the changing landscape of therapeutic approaches is helpful in assessing strategic patterns in the approach to therapy, but the information is scattered throughout the literature and public data resources.

**Results:** We gathered data on therapeutic strategies for 299 diseases into the Drug Database for Inborn Errors of Metabolism (DDIEM). Therapeutic approaches, including both successful and ineffective treatments, were manually classified by their mechanisms of action using a new ontology.

**Conclusions:** We present a manually curated, ontologically formalized knowledgebase of drugs, therapeutic procedures, and mitigated phenotypes. DDIEM is freely available through a web interface and for download at http://ddiem.phenomebrowser.net.

## Background

Rare hereditary diseases together make up a significant component of overall morbidity and mortality around the world. A disease is considered rare in the United States (US) if it affects fewer than 200,000 individuals. With more than 7,000 rare diseases characterized, the overall population burden is therefore of the order of 20 to 30 million [1]. Consequently, it has become a significant concern for strategic health funding bodies in North America, Europe, and elsewhere to coordinate international infrastructure, research and development, and delivery of new therapeutics [2]. While much effort has gone into mobilizing patient data across national boundaries and undertaking extensive genetic studies, the Orphan drug initiatives in the US starting in 1983, together with ongoing programs at the FDA and NIH, as well as those funded by the European Commission and Canadian authorities, saw a dramatic increase in therapies for rare diseases. These were notable for a large number of drug repurposing successes, providing accelerated access to patients [3, 4, 5]. Currently, around 15% of new orphan drug approvals are for metabolic and endocrine therapies, most of these through the application of small molecules [6].

Metabolic hereditary diseases, or inborn errors of metabolism (IEMs), are a group of disorders that disrupt normal metabolism and physiology, ultimately affecting almost all biochemical pathways and processes in the body. Individually metabolic diseases are rare, but, as with rare disease as a whole, relatively common when considered as a class of disease [7]. Over 1,000 distinct hereditary metabolic diseases have been identified and recently assigned to 130 categories in a new nosology [8], although the underlying causes are only understood for about half of these diseases.

Data resources have been developed for hereditary metabolic diseases. For example, the Rare Metabolic Diseases (RAMEDIS) database [9] contains information on rare diseases and their treatment options. However, RAMEDIS covers only 93 different IEMs and primarily relies on information from case reports and their associated available data on treatment options for most of the 93 disorders covered. IEMbase is a database that contains clinical, biochemical, and genetic information on 1,310 IEMs, as well as a nosology that classifies each disease into one of ten significant categories [10]. However, neither RAMEDIS nor IEMbase cover different treatment options and strategies for rare metabolic diseases.

Therapeutic approaches to inherited metabolic diseases are diverse and have to distinguish, for example, between loss of function, dominant gain of function, and the generation of toxic metabolic intermediates. Among therapeutic strategies that address metabolic diseases are gene therapy, metabolite level manipulation, transplantation or surgery, and small molecule therapies to stabilize or enhance residual enzyme activity. Therapies may address the underlying mechanism directly or indirectly. They may provide symptomatic or prophylactic therapy where some specific phenotypes of the syndrome are addressed, and sometimes they improve the long term pathological outcomes [11, 12, 13]. More importantly, a large number of rare IEMs do not currently have an approved treatment approach, and new therapeutic strategies are tested in individual cases or clinical trials, and are subsequently reported in the scientific literature [14]. An overview of the strategic landscape of therapeutic approaches to treating IEMs is of potential use not only for investigators developing new therapies but also for regulators and research funders.

We have developed the Drug Database for Inborn Errors of Metabolism (DDIEM), a database that covers experimental approaches, clinical treatments (established as well as investigational therapies) and their outcome, for diagnosed rare metabolic diseases. Individual diseases are linked to established information resources and databases. In order to classify therapeutic approaches, we have created a new ontology of strategies for treating IEMs and categorized treatments using this ontology. The use of an ontology allows for data integration, aggregation, and query expansion, as well as enhancing data access according to the FAIR (Findable, Accessible, Interoperable, Reusable) data principles [15]. We used a wide variety of public online data sources and, where possible, we extracted the specific aspects of phenotype addressed by the therapy from the literature through manual expert curation. DDIEM is freely available at http://ddiem.phenomebrowser.net.

## Results

DDIEM is a database of therapeutic approaches to treating IEMs which have been manually extracted from the scientific literature. DDIEM currently covers 299 rare diseases along with the association of 305 genes and 552 drugs that were used to treat these diseases; these treatment attempts have been linked to 1,455 distinct disease-associated phenotypes that were influenced by these treatments. The main entity in DDIEM is the therapeutic procedure which is used to treat a metabolic disorder; these procedures are classified based on their underlying mechanism or modality. We have developed an ontology using the Web Ontology Language (OWL) [16] to formalize this classification of treatments in DDIEM.

### Classification of treatment mechanisms: the DDIEM ontology

There are several existing strategies for classifying treatments for inborn errors of metabolism. Most of these are concerned with the underlying biochemical error and affected pathways. For example, ICD-11, based on the recommendations of the Society for the Study of Inborn Errors of Metabolism (SSIEM) [17], classifies in-born errors of metabolism mainly by type of metabolism; e.g., lipid metabolism, carbohydrate metabolism. ICD-11 also has other general axes of classification for IEMs, such as peroxisomal diseases, and more specific axes of classification, such as classifying diseases based on involvement in purine metabolism. Other nosologies emphasize diagnostic and clinical aspects of disease and, for example, classify diseases into disorders that either involve only one functional system (such as the immune system) or diseases in which the basic biochemical lesion affects metabolic pathways common to a large number of tissues [17]. The latter group is divided by molecule type of interest and energy metabolism. More recently, a nosology of IEM has been developed which classifies diseases by biochemical entity and process, a work that emerged from the European Reference Network for Hereditary Metabolic Disorders [8]. At least one classification is based neither on biochemical pathway nor clinical manifestation, but rather on modalities of treatment [12, 18].

Our major interest concerns medical treatment rather than dietary modification (i.e., excluding or augmenting the diet with naturally occurring food substances). However, most of the available therapeutic options for IEMs are symptomatic where they treat the symptoms or downstream effects without addressing the direct effects of the altered protein underlying the disease. Some other approaches include Enzyme Replacement Therapy (ERT) where the missing enzyme is replaced by infusions of an enzyme that is purified from human or animal tissue or blood or produced by novel recombinant techniques [19]. Typically, the enzyme is modified to allow for a longer half-life, more potent activity, resistance to degradation or targeting to a specific organ, tissue or cell type.

Some treatment strategies deliver their effects only if there is some residual enzymic activity and therefore their efficacy is dependent on this. Examples are substrate reduction therapy (SRT) [11], chemical chaperone therapy [20] and pharmacological chaperone therapy [21]. SRT is a strategy that works through limiting the amount of substrate synthesized to a level that can be effectively cleared by the impaired enzyme. The efficacy of SRT is mutation-specific and dependent on residual enzyme activity level [22]. Chaperone therapy uses small molecule substances, often osmolytes. These either facilitate folding or stabilize misfolded proteins to rescue residual activity, for example by reducing abnormal aggregation or interacting with active sites. [20]. In some cases a disease is amenable to direct gene therapy [23] which usually alters the somatic genome to prevent or treat a disease through insertion of a functional copy of the affected gene [20]. More recently, the potential for gene editing and epigenetic modifications are showing considerable promise [24].

We have developed and implemented the DDIEM ontology as a framework for classifying treatments by mechanism of action, as described above. This ontology allows the generation of a well-structured dataset broadly interoperable with existing resources. We based the DDIEM ontology on the Ontology of General Medical Sciences (OGMS) [25] and have made it available in the community ontology repositories BioPortal [26] and AberOWL [27].

In the DDIEM ontology, we classify mechanisms of treatment using three upper level classes. The first class covers those treatments that attempt to compensate for or modulate the biological functions affected by the dysfunctional protein (`mechanistically predicated therapeutic procedure`). The second class covers those that treat symptoms (`symptomatic therapeutic procedure`); and the third class covers surgical or physical procedures such as stem cell transplantation (`surgical or physical therapeutic procedure`). Some therapeutic strategies combine multiple drugs or other treatment modalities which work through different mechanisms, for example one drug addressing the primary lesion and another its symptomatic consequences. These combination therapies are flagged using the ontology. Figure 1 depicts the structure of the DDIEM ontology; the definition of each class can be found in Table 1. Table 2 shows several examples of therapeutic procedure for the main classes in DDIEM.

**Table 1:** A summary of the main mechanisms of therapeutic procedures for metabolic diseases.

| Therapeutic procedure | Definition |
| --- | --- |
| Combination therapeutic procedure | A therapeutic procedure using multiple therapies to treat a single disease. Often all the therapies are pharmaceutical. It can also involve non-drug therapy, such as the combination of medications, behaviour modification, or physical procedures. The different components might act through the same or different mechanisms. |
| Mechanistically predicated therapeutic procedure | A therapeutic procedure which directly addresses the effects of a protein defect produced by the mutation, its structure, activity, or end product. Therapies which target the local interaction or regulatory network or pathway in which the affected protein lies are also regarded as mechanistic therapies. |
| Complementation therapy | A therapeutic procedure in which a treatment directly or indirectly compensates for the loss or gain of activity of a genetically defective protein within the network or pathway of which that protein is a member. |
| Direct complementation of a genetically defective protein | A therapeutic procedure in which a genetically defective protein is replaced by a canonical source of the same protein, genetically as in gene therapy, or by some other means of delivery, whose completion is hypothesized by a health care provider to eliminate a disorder or to alleviate the signs and symptoms of a disorder or pathological process. |
| Compensatory complementation of a genetically defective protein | A therapeutic procedure in which the availability, activity, stability, or turnover of a defective enzyme is modified by delivery of small or macro-molecules, or genetic or epigenetic manipulation. |
| Functional complementation of a genetically defective protein | A therapeutic procedure in which the composition or metabolic activity of the pathway or network in which the defective protein is found is modified, compensating for alteration of activity of that protein. |
| Dietary regime modification | A therapeutic procedure in which the diet is supplemented with or depleted in molecules closely related to the products or end processes of a genetically defective protein which occur as natural products in the diet. |
| Dietary exclusion | A therapeutic procedure in which the diet is depleted in molecules closely related to the products or end processes of a genetically defective protein, which occur naturally in the diet. |
| Dietary supplementation | A therapeutic procedure in which the diet is supplemented with foodstuffs containing molecules closely related to the products or end processes of a genetically defective protein, which occur naturally in the diet. |
| Metabolite replacement | A therapeutic procedure involving enteral, parenteral, or transdermal provision of small molecules closely related to the products or end processes of a genetically defective protein. |
| Activity modification of a genetically defective protein | A therapeutic procedure in which generally small molecules are delivered to the organism in order to directly increase, decrease, or alter the activity or stability of a genetically defective protein. |
| Symptomatic therapeutic procedure | A therapeutic procedure aimed at amelioration of one or more abnormal phenotypes generated as a consequence of a defective protein or process by a means unrelated to the immediate pathway or network environment of the defective protein. Generally working at the level of the tissue or overall organismal physiology. |
| Surgical or physical therapeutic procedure | A therapeutic procedure to mitigate the immediate or future effects of the presence of a genetically defective protein which involves physical conditioning or anatomical modification. |
| Functional complementation of a genetically defective protein by inhibition | A therapeutic procedure that inhibits a component of the pathway or network in which a genetically defective protein is found, compensating for alteration of activity of that protein. |
| Functional complementation of a genetically defective protein by stimulation | A therapeutic procedure that stimulates a component of the pathway or network in which a genetically defective protein is found, compensating for alteration of activity of that protein. |

**Table 2:** Examples of the main mechanisms of therapeutic procedures for metabolic diseases.

| Therapeutic procedure | Examples |
| --- | --- |
| Activity modification of a genetically defective protein | In Bartter syndrome type 4A (OMIM:602522), tanesipimycin (17-allylamino-17-demethoxygeldanamycin or 17-AAG), an Hsp90 inhibitor, enhances the plasma membrane expression of mutant barttins (R8L and G47R) in Madin–Darby canine kidney cells [28]. |
| Activity modification of a genetically defective protein by epigenetic manipulation | HDACi 109/RG2833 increases FXN mRNA levels and frataxin protein, with concomitant changes in the epigenetic state of the gene, in the treatment of Friedreich's ataxia (OMIM:229300) [29]. |
| Activity modification of a genetically defective protein by genome editing | CRISPR-Cas9-mediated gene editing is a potential treatment for Hurler syndrome (OMIM:607014) as established in cell and animal based studies [30] [31]. |
| Activity modification of a genetically defective protein by transcriptional or translational modification | Vitamin D in treating hyperprolinemia (OMIM:239500) and recombinant human erythropoietin (rhuEPO) in treating Friedreich's ataxia (OMIM:229300) are examples of drugs that enhance the activity of PRODH and FXN respectively by transcriptional modulation of PRODH in hyperprolinemia and increasing frataxin expression in Friedreich's ataxia [32] [33]. |
| Direct complementation of a genetically defective protein | Recombinant human IGF1 (mecasermin) is a form of enzyme replacement therapy in treating insulin-like growth factor I deficiency (OMIM:608747) [34]. |
| Direct complementation of a genetically defective protein by gene therapy | An adeno-associated viral vector containing a porphobilinogen deaminase gene is a treatment for acute intermittent porphyria (OMIM:176000) [35]. |
| Functional complementation of a genetically defective protein by inhibition | Miglustat is a form of substrate reduction therapy (SRT) used to treat Gaucher's disease (OMIM:230800) where miglustat inhibits the ceramide-specific glucosyltransferase which catalyzes the first committed step of GSL synthesis [36]. |
| Procedure to mitigate dominant effect of a genetically abnormal protein | In treating hyperinsulinism-hyperammonemia syndrome (OMIM:606762), diazoxide inhibits insulin release and promotes complete resolution of hypoglycemia [37]. |
| Dietary exclusion | A valine-restricted diet is a treatment for patients suffering from isobutyryl-CoA dehydrogenase deficiency (OMIM:611283), which involves valine metabolism [38]. |
| Dietary supplementation | Beneprotein and uncooked cornstarch as high sources of protein and carbohydrates are used to prevent long-term complications in glycogen storage disease IXb (OMIM:261750) patients [39]. |
| Metabolite replacement | Hydrocortisone as a replacement therapy for patients with 17-alpha-hydroxylase deficiency (OMIM:202110) [40]. |
| Functional complementation of a genetically defective protein by stimulation | Sodium valproate activates the expression of one glycogen phosphorylase isoform, GP-BB, which in turn results in a decrease in intracellular glycogen accumulation, a dominant feature of glycogen storage disease type V (OMIM:232600) [41]. |

**Figure 1:**
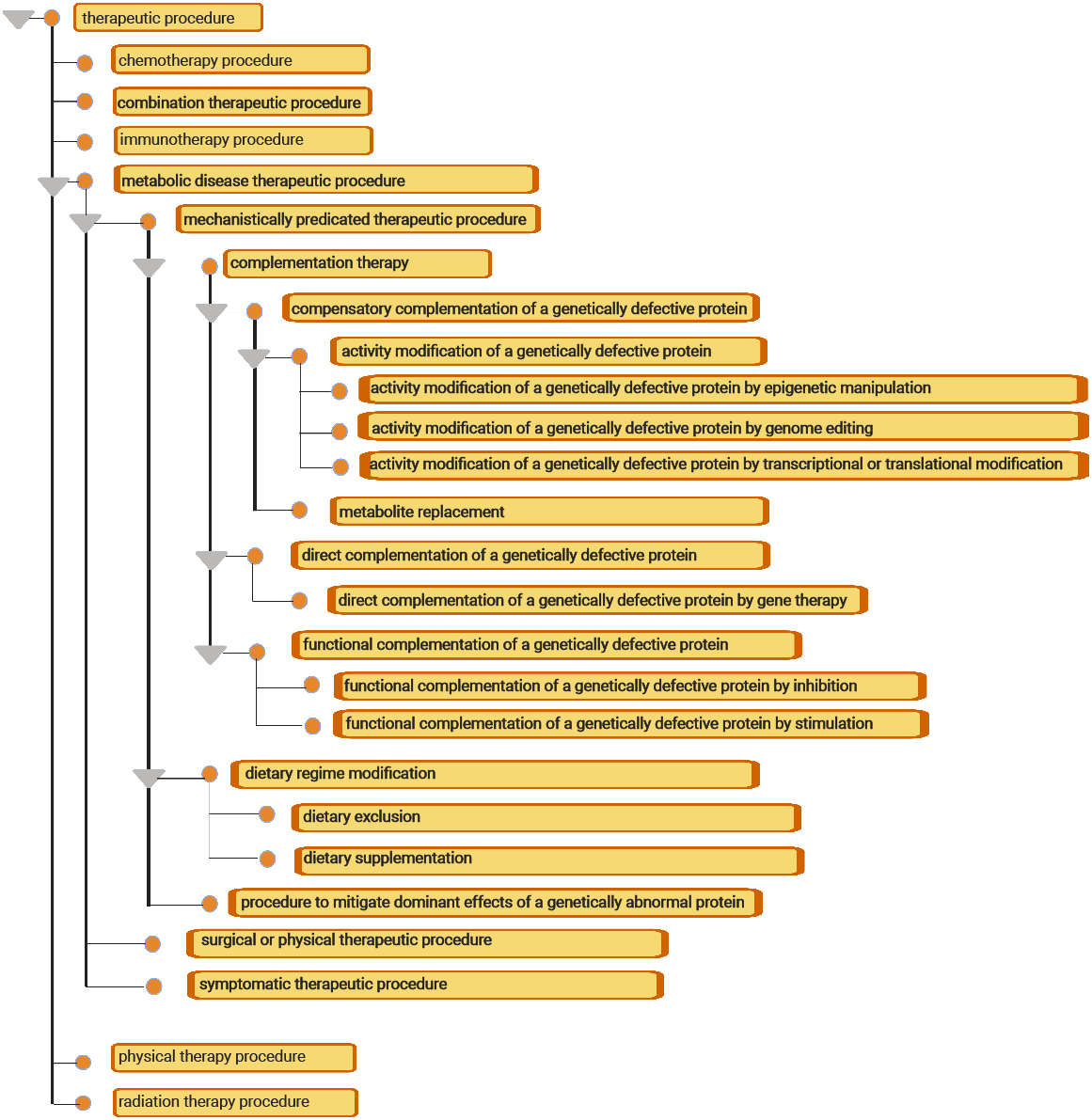
Structure of the therapeutic ontology

The DDIEM ontology is based on an upper-level ontology for medical sciences, the OGMS. Reusing such an upper-level ontology allows us to consistently integrate the DDIEM ontology with related efforts of formalizing categories in the biomedical domain, including the Drug Ontology [42] and the Medical Action Ontology [43] which is currently under development and will broadly characterize therapeutic procedures.

### DDIEM database

In DDIEM, we manually curated over 1,600 scientific literature articles to record evidence for therapies that have been attempted for currently 299 rare metabolic diseases; these 299 metabolic diseases are associated with 305 genes. We classify therapeutic approaches that have been reported in literature for each disease in DDIEM using the DDIEM ontology. The distribution of the drugs used as part of therapeutic procedures and their classification using the DDIEM ontology is shown in Figure 2. These therapeutic approaches involve 552 unique drugs and correct or ameliorate 1,455 distinct phenotypes. For each therapeutic strategy, we record both evidence and provenance information using standardized identifiers and coding systems. Figure 3 provides an overview of the type of information we collect and the relations between the different types of information.

**Figure 2:**
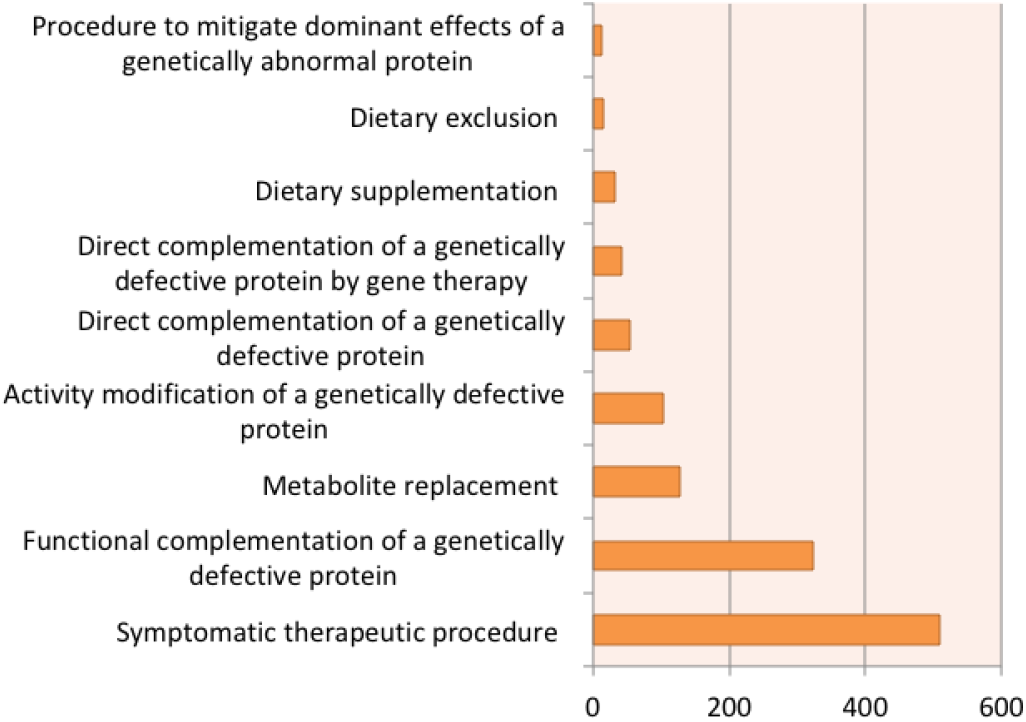
Distribution of the Drugs based on classes in the DDIEM ontology

**Figure 3:**
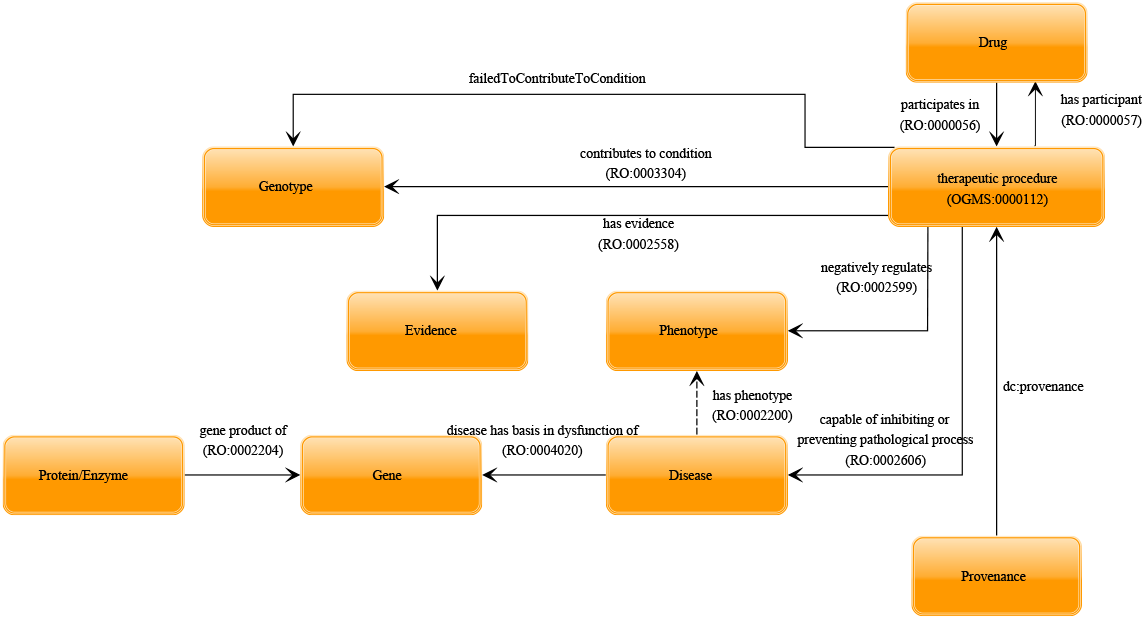
Overview of entity types and their relations in DDIEM.

We identified 30 diseases where the outcomes of a therapeutic procedure were affected by particular genotypes or genetic variants. Modifying genotypes may include genotypes that affect the pharmacological action of a substance, either positively or negatively. For example, in familial hyperinsulinemic hypoglycemia type 3 (`OMIM:602485`), the Y214C variant in the *GCK* gene was found to result in patients’ being unresponsive to diazoxide while patients carrying the M197I variant in the same gene did respond to the treatment [44]. When available, we record this information in DDIEM.

There are 55 orphan diseases in DDIEM for which we did not assign any therapeutic strategy class, either due to absence of information on tested therapeutic interventions in the literature, or the benign status of the disease which does not usually require any intervention. In the situation where we were unable to identify any report of a therapeutic strategy for a disease in DDIEM, we mark the disease as “no treatment is available”.

Phenotypes in DDIEM that are affected by a therapeutic procedure are formally coded using the Human Phenotype Ontology (HPO) [45] or, if no HPO class could be identified, using the Mammalian Phenotype Ontology (MPO) [46]. In 113 cases we could not identify a phenotype class matching the described phenotype in these two ontologies and recorded the phenotype as free text; additionally, we requested extension of the HPO with the missing phenotypes so that we can formally include them in DDIEM once they become available in the HPO.

In DDIEM, we distinguish between six different types of evidence provided for the affect of a therapeutic procedure on disease-associated phenotypes, and we use the Evidence and Conclusion Ontology [47] to record evidence for our assertions. We distinguish evidence based on animal models (`ECO:0000179`), clinical trials `ECO:0007121`), and experiments on cell lines (`ECO:0001565`). In some cases, study authors suggested therapeutic procedures based on clinical observation alone; we record these as inference by a study author (`ECO:0007764`). In rare cases, DDIEM curators inferred information about a therapeutic procedure although it was not explicitly stated in the article; we mark this using the “inferred by curator” evidence (`ECO:0000305`). Figure 4 shows the distribution of the evidence codes in DDIEM.

**Figure 4:**
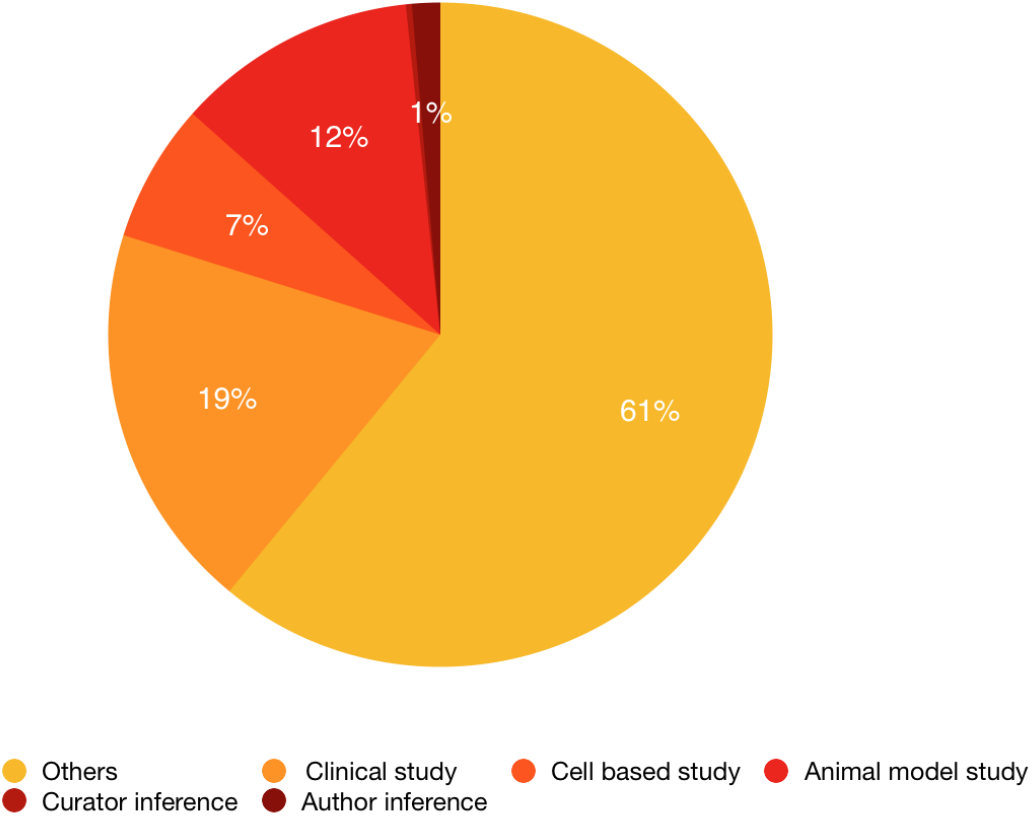
Evidence code distribution in DDIEM.

### Implementation of FAIR principles

DDIEM is intended as a resource for biomedical and clinical researchers as well as for computational scientists. To enable DDIEM content to be usable by a wide range of researchers, we aim to follow the FAIR principles (Findable, Accessible, Interoperable, and Reusable) [15].

Specifically, to ensure interoperability, we followed the Linked Data principles [48] and linked the entities described in DDIEM to community reference resources and ontologies. Diseases are mapped to Online Mendelian Inheritance in Man (OMIM) [49] disease identifiers as well as to identifiers in the IEMbase database [10]. Genes are linked to the NCBI Entrez gene database [50] and their products to Expasy [51], KEGG [52], and the UniProt [53] databases. We mapped drugs to Drugbank [54], PubChem [55], ChEBI [56], and to identifiers from the Anatomical Therapeutic Chemical Classification System (ATC) [57]. In DDIEM, we rely on ontologies in the OBO Foundry [58] as collaboratively developed reference ontologies in the biomedical domain. We represent phenotypes using either the Human Phenotype Ontology (HPO) [45] or the Mammalian Phenotype Ontology (MPO) [46], and we use the Evidence and Conclusion Ontology (ECO) [59] to specify different study types and evidences.

DDIEM content is accessible through a website as well as through a public SPARQL end-point to enable computational access. DDIEM data is also downloadable, and each release of the data is assigned a unique Digital Object Identifier (DOI) [60]. To make DDIEM content findable, we registered DDIEM on the FAIR-sharing platform [61] and the DDIEM ontology in several ontology repositories.

## Discussion

There exist several formal data resources for information on therapies for rare metabolic diseases. However, the majority of them focus on symptoms, clinical or metabolic aspects rather than therapeutic approaches. For example, the Orphanet [14] database includes drugs designated for treating rare genetic diseases. Similarly, RAMEDIS [9] focuses on treatments described in case reports for a number of metabolic diseases. However, neither provide information on the mechanism of drug mechanism or on the phenotypes corrected or alleviated. Another database, IEMbase [10], provides some therapeutic information but concentrates mainly on the clinical and biochemical aspects of rare metabolic disease, and provides an expert platform to facilitate their early and accurate diagnosis. To the best of our knowledge, DDIEM is the first database that significantly focuses on the treatment either used or investigated, to correct or alleviate phenotypes associated with the course of a specific metabolic disease.

With a focus on phenotypes and mechanisms, DDIEM uniquely addresses fundamental aspects of rare disease therapy with the aim of supporting the analysis of the therapeutic landscape and the development of new treatments. The Data Mining and Repurposing (DMR) task force of the IRDiRC [62] recently emphasized the important contribution of data mining and data integration to the development of new drugs for rare diseases, and highlighted the need to get data out of silos – often semantic silos – to facilitate *in silico* approaches to drug development and repurposing. DDIEM is developed on the FAIR principles to ensure findability, accessibility, interoperability, and reusability and is accessible either through a web interface or computationally through the DDIEM API. Use of community standards, databases, and ontologies permits ready computational integration of curated DDIEM information into other datasets. DDIEM therefore provides a dataset that can be reused by a wide range of researchers, using different methodological approaches, to investigate existing and develop new therapeutic approaches.

The current data in DDIEM does not cover the details or scale of clinical trials or individual reports, the drug formulation, or the dosage applied. Such information could further provide valuable information particularly for drug repurposing. In the future, we plan to expand DDIEM by adding a limited amount of quantitative information and dosage, as well as reporting quantitative drug effects.

## Conclusion

We developed DDIEM, a database focusing on treatments for Inborn Errors of Metabolism. DDIEM integrates literature-reported treatments for a wide range of IEMs together with information on the outcome. The reports range from individual case reports to clinical trials. Consequently, DDIEM provides information on the on- and off-label uses of drugs and their effects for treating IEMs and may be used to accelerate drug development and drug repurposing for these diseases.

In creating DDIEM we have extensively analyzed publicly available data on the treatment of rare metabolic diseases, and present it in a way which permits integration with other datasets and resources. The value we add to the currently available public data is in our expert curation and semantic formalization, integrating and presenting this data in a way compliant with FAIR standards, and making it freely accessible to investigators in both public and private domains. DDIEM is accessible through its website and can be queried computationally. We support the conclusions and recommendations of the IRDiRC regarding data integration and exploitation [62], and believe that DDIEM will contribute to the discovery or repurposing of drugs for diseases still lacking effective therapies.

## Materials and Methods

### Resources used

We rely on literature resources listed in PubMed and information published on ClinicalTrials.gov to gather information about treatments that were used for rare metabolic diseases. For parts of the curation process we used the GoPubMed software [63] to access and query literature resources. While PubMed reports on rather complete studies, ClinicalTrials.gov provides information for ongoing studies with potential interim results for each rare disease. We obtained a list of metabolic diseases from the Genetic and Rare Diseases (GARD) information center database [64].

### Literature curation

To create the DDIEM content, we searched in literature the disease names or synonyms from the metabolic disease list we gathered from GARD. In the cases where we could not find any literature records for the name or synonyms of a given disease, we used the gene name associated with the disease in the search. We then filtered all the retrieved records by selecting the ones containing treatment or management information. Since these diseases are considered rare, we did not exclude any literature as long as it provides information about therapeutic interventions, and in which we found explicit evidence for the correlation between the applied treatment and clinical phenotypes, which are either completely-treated or alleviated. However, in DDIEM, we focused mainly on pharmacological and, to some extent, on dietary interventions. We then curated the resulting literature documents manually to extract the metabolic disorder investigated, the drugs used to treat the disorder, the effects of the treatment, and the suggested mechanism of action. If drugs exerted their impact differently between patients with the same metabolic disease due to different genetic variants in the same gene, we added those variants as an additional reference. We categorized variants that affect the effectiveness and success of therapeutic intervention into two categories based on whether they promoted the drug effect or not.

We provided all literature references as provenance information through the DDIEM web interface. After mapping most drugs to their identifiers in the cross-linked databases, there were some therapies we could not link to any database. Hence, we indicated their identifiers as non-available (NA).

Additionally, we developed a therapeutic ontology that describes the mechanism by which the therapeutic procedures act to either adequately treat the disease or modify disease pathogenesis. We recorded the disease phenotypes which were treated or improved by the therapeutic procedure and mapped them to reference ontologies. Similarly, where phenotypes had no matching identifiers in those ontologies, we referred to identifiers as non-available (NA).

### Data representation and web interface

We represent the content of DDIEM using the Resource Description Framework (RDF) [65]. First, we normalized the curated data (which is primarily kept in a tab-separated form) by mapping the DDIEM content to their identifiers in different databases (Uniprot, Expasy, KEGG, OMIM and Drugbank). In the second step, we represent the DDIEM content using the DDIEM RDF data model which captures the relations between the biomedical entitites we characterize in DDIEM.

We developed a web interface to allow users to navigate the DDIEM database. The interface provides a list of diseases covered in the database along with their details including treatments, participating drugs, phenotypes, and references to the publications from which the information was extracted. The web interface also provides a search in the DDIEM resources using disease, drug, and mode of action of the therapeutic procedure. Furthermore, to enable automated access to the DDIEM content, we provide a SPARQL endpoint which directly queries the RDF data we provide.

We further implemented a web server using Node.js and developed an API for accessing DDIEM resources. The API response is formatted in either JSON or JSON-LD format which enables the possibility to build client applications using the RDF data underlying DDIEM. We used OpenLink’s Virtuoso database as an RDF store to store and query the DDIEM data, and all primary data is stored in the RDF store and the Node.js server retrieves all data through SPARQL queries.

The DDIEM data is also available for download in RDF format from the DDIEM website at http://ddiem.phenomebrowser.net.

## Abbreviations

ChEBI: Chemical Entities of Biological Interest
DDIEM: Drug database of inborn errors of metabolism
DMR: Data Mining and Repurposing
ECO: Evidence and Conclusion Ontology
ERT: Enzyme replacement therapy
FAIR: Findable, Accessible, Interoperable, Reusable
FDA: Food and Drug Administration
GARD: Genetic and Rare Diseases Information Center
HPO: Human Phenotype Ontology
ICD: International Classification of Diseases
IEM: Inborn errors of metabolism
IRDiRC: International Rare Diseases Research Consortium
KEGG: Kyoto Encyclopedia of Genes and Genomes
MAxO: Medical Action Ontology
MPO: Mammalian Phenotype Ontology
NA: Not available
NCBI: National Center for Biotechnology Information
NIH: National Institutes of Health
OGMS: Ontology for General Medical Science
OMIM: Online Mendelian Inheritance in Man
OWL: Web Ontology Language
RAMEDIS: Rare Metabolic Diseases Database
RDF: Resource Description Framework
SRT: Substrate reduction therapy
SSIEM: Society for the Study of Inborn Errors of Metabolism
WHO: World Health Organization

## Ethics approval and consent to participate

Not applicable.

## Consent for publication

Not applicable.

## Availability of data and materials

DDIEM is freely available at http://ddiem.phenomebrowser.net. Source code and all data used to construct DDIEM is also freely available on https://github.com/bio-ontology-research-group/DDIEM.

## Author’s contributions

RH conceived of, and PNS and RH designed the study. MA and EM performed the literature curation and creation of content. ARS and AAK implemented the DDIEM database and website and generated the RDF datasets. SK provided technical expertise on literature curation and database design. MA, EM, PNS, RH, and ARS developed the DDIEM ontology. SK, PNS, RH supervised the work. MA, SK, PNS, and RH drafted the manuscript, and all authors critically revised the manuscript for scientific content and accuracy. All authors have read and approved the final manuscript.

## Acknowledgements

The authors acknowledge the support and generous assistance of Prof. Wyeth W. Wasserman, Prof. Nenad Blau and Tamar Av-Shalom in facilitating linkage between DDIEM and IEMbase. We thank Prof Michel Dumontier for advise on the curation process, data representation, and implementation of FAIR principles.

